# Evidence for current circulation of an ancient West Nile virus strain (NY99) in Brazil

**DOI:** 10.1101/2020.09.21.307199

**Authors:** Márcio Junio Lima Siconelli, Daniel Macedo de Melo Jorge, Luiza Antunes de Castro-Jorge, Antônio Augusto Fonseca-Júnior, Mateus Laguardia Nascimento, Vitor Gonçalves Floriano, Fransérgio Rocha de Souza, Eudson Maia de Queiroz-Júnior, Marcelo Fernandes Camargos, Eliana Dea Lara Costa, Adolorata Aparecida Bianco Carvalho, Benedito Antonio Lopes da Fonseca

## Abstract

**Introduction:** In Brazil, West Nile virus (WNV) was first detected in 2018 from horses with neurological disease.

**Aim:** Here we present the first reported case in Ceará state and complete genome sequence from an isolated in Espírito Santo state from 2019.

**Methods:** The virus was isolated from a horse that died with neurological signs in Espírito Santo and was sequenced by MiSeq.

**Results:** Phylogenetic analysis reveals that this isolate belongs to lineage 1a, clustering with NY99 strain, that disappeared from the USA since 2005.

**Conclusions:** Our findings reinforce the hypothesis that WNV has been silently circulating in Brazil for many years.

---

West Nile virus (WNV) is a zoonotic RNA virus from the Flaviviridae family and Flavivirus genus. It belongs to the Japanese Encephalitis virus serocomplex that contains viruses (e.g. Usutu virus, Saint Louis virus, Murray Valley Encephalitis virus and Cacipacore virus) that cause neurological disease in humans and animals^1,2^. *Culex* mosquitoes are the main vectors for enzootic or epizootic WNV transmission; birds are the amplifier hosts, and big mammals, such as horses, accidental hosts^3^. In the Americas, WNV was first detected in New York City in 1999, probably introduced from Israel^4^, and since then spread to many other American countries. Almost 20 years later, in 2018, the first molecular detection and viral isolation in Brazil was accomplished from samples collected from a horse with neurological signs in a locality from Espírito Santo state^5^. Before that, there was serological evidence of WNV circulation in horses in Paraíba state^3^, but only in 2014, the first official human case was reported in Piauí state^6^.

In June 2019, two fatal cases in horses with neurological disease were reported, one from the municipality of João Neiva, in Espírito Santo state (804_02_ES), and another from Boa Viagem, in Ceará (827_01_CE) **(Figure 1)**. Clinical signs included neurological and locomotor disorders as muscle tremors/rigidity, shaking head, weakness, ataxia, recumbency, hyperesthesia, limbs paresis, pedaling movements, and mydriasis. Central Nervous System (CNS) samples were collected for diagnosis.

**Figure 1.**
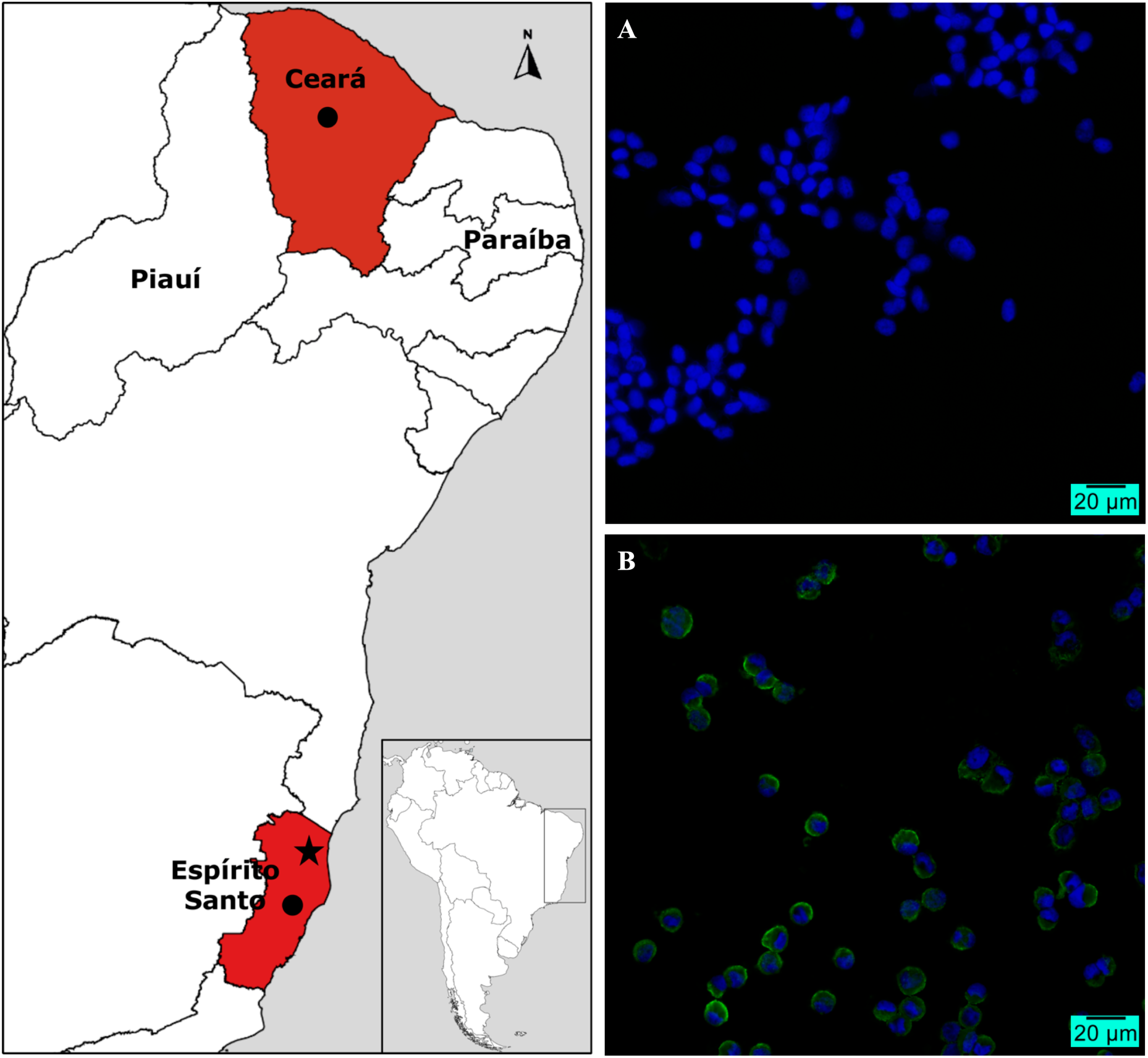
Map of states (red) with molecular evidence of WNV circulation in horses with neurological disease. Black dots: municipalities of occurrence of WNV, João Neiva in Espírito Santo (ES), and Boa Viagem in Ceará; Black star: municipality of São Mateus/ES where the first isolation/complete sequencing was done in 2018; Indirect immunofluorescence assay for detection of West Nile virus antigens in VERO cells in magnification of 40X. (A) Mock cells (uninfected cells as negative control); (B) Infected cells from the 2nd passage of WNV isolated after 48h post-inoculation. Hyperimmune serum obtained from infected mice with WNV reference NY99 strain was used in both mock and infected cells. Blue: nucleus stained with DAPI; Green: WNV antigen.

As expected, rabies investigation resulted negative since both animals both had been recently vaccinated. For further diagnosis investigation, samples were sent to the Federal Agricultural Defense Laboratory of Minas Gerais (LFDA/MG) from the Ministry of Agriculture, Livestock, and Supply (MAPA). The animals tested positive for WNV by the World Organization for Animal Health (OIE) RT-qPCR protocol^7,8^. CNS frozen samples were then sent to the Molecular Virology Laboratory of Ribeirão Preto Medical School – University of São Paulo (LVM-FMRP-USP) for isolation and genome characterization.

The samples were thawed on ice and 0.03-0.05 grams of spinal cord (804_02_ES) and encephalon (827_01_CE) were manually macerated with a microcentrifuge pestle (Corning^®^) in DMEM (Hyclone™) supplemented with 2% fetal bovine serum (FBS) and 2% antibiotic-antimycotic (Cultilab™). The homogenate was clarified at 16,000 g for 10 minutes at 4°C and the supernatant were inoculated into VERO (CCL 81) and C6/36 (CRL 1660) cells for 1h to adsorption. Then, maintenance medium (2% FBS) was added and cells incubated at 37°C (VERO) and 28°C (C6/36). After 5-6 days cytopathic effect was clearly observed in VERO cells and WNV was successfully isolated from 804_02_ES in both cell types and a second inoculation on both cells was done to increase virus titer.

Virus isolation was confirmed by RT-qPCR^7,8^ and indirect immunofluorescence assay (IFA) using hyperimmune serum from infected mice with WNV NY99 reference strain (Figure 1). The isolate (804_02_ES) and a clinical sample from Ceará state (827_01_CE) were sequenced by MiSeq Illumina sequencing platform (https://www.illumina.com) on the Life Sciences Core Facility (LMSeq) of São Paulo State University (UNESP/FCAV).

We conducted *de novo* analysis using a bioinformatic pipeline focused on viruses. The softwares used on the analysis were FastQC version 0.11.8 (https://www.bioinformatics.babraham.ac.uk/projects/fastqc), Trimmomatic version 0.3.9 (http://www.usadellab.org/cms/?page=trimmomatic) and AfterQC version 0.9.7 (https://github.com/OpenGene/AfterQC). The virus assembly was mapped against NY99 strain (GenBank accession: MH643887) in order to obtain a consensus sequence. The final consensus sequence of 804_02_ES was 10,893 nucleotides in length with an identity of 99.71% and 93.44%, for nucleotide and protein, respectively.

The 804_02_ES genome was aligned with MAFFT 7.0 software using a dataset obtained from NCBI and ViPR (https://www.viprbrc.org). A total of 116 sequences (1953– 2018) from worldwide distribution were selected. The dataset was edited and curated to use in the phylogenetic tree to keep the entire ORF. The Maximum-Likelihood (ML) tree was performed using IQ-TREE 2.0 software (http://www.iqtree.org) in automatic selection, being GTR+F+I+G4 the best nucleotide substitution model, with support analysis of 10,000 replicates. The tree graph was edited with iTOL v.5 (https://itol.embl.de).

The 804_02_ES (GenBank accession: MT905060) genome was phylogenetically determined to belong to WNV lineage 1a **(Figure 2)** in accordance with the first isolation occurred in 2018 (GenBank accession: MH643887), also from Espírito Santo state (municipality of São Mateus) a region nearby the place where the sample yielding this isolate was collected and diverging in only 32 bases^5^. TempEst v1.5.3 was used to perform the molecular clock to find the time of the Most Recent Common Ancestors (tMRCA), and to inspect and identify any inconsistency in our sequence according to all databases of temporal structures.

**Figure 2.**
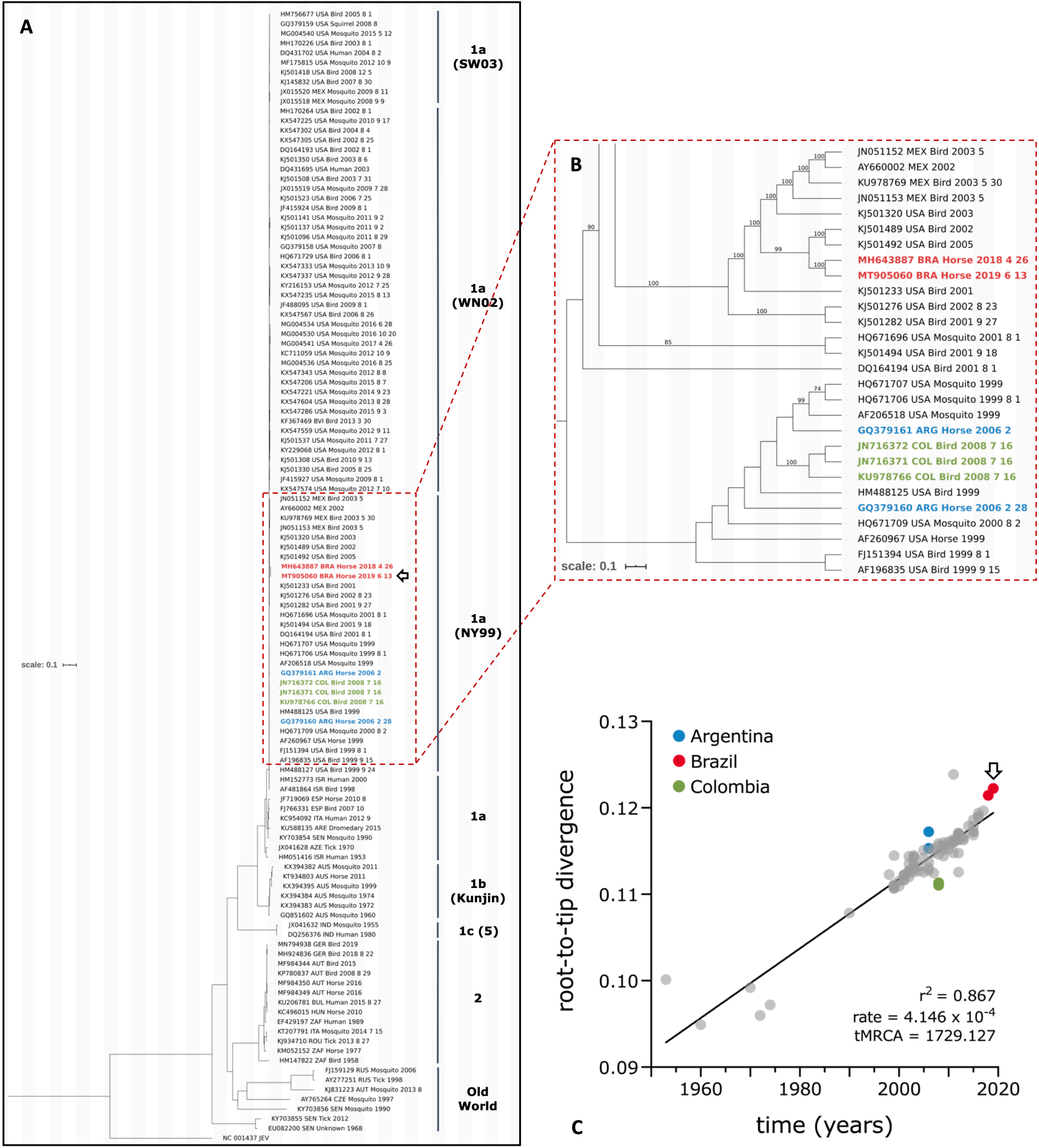
Maximum-likelihood (ML) phylogenetic reconstruction of WNV from 1953–2019. (A) 117 selected complete sequences from the global dataset were included in this analysis, presenting 0.1 substitutions per site. The GTR+F+I+G4 substitution model and support analysis of 10,000 replicates were used to obtain the phylogenetic tree; (B) dashed box highlighting South America (SA) strains, including the new isolate (MT905060), that cluster with lineage 1a NY99 strain; only bootstrap values equal or bigger than 70% were shown; (C) root-to-tip analyses obtained with TempEst, using ML reconstruction obtained of 94 complete genomes from 1a and 1b lineages; graph correlation between time (years) and genetic divergence (substitutions per site) from the root of the tree (time to the Most Recent Common Ancestors, tMRCA) to the tips (sampled genomes) is being shown; evolutionary slope rate: 4.146 x 10^−4^ substitutions/site/year; black arrow outline = new isolate. All colored strains belong to SA: Brazil (red), Argentina (blue), and Colombia (green). Old world lineages include: 3 (Rabensburg), 4a, 4b (9), 7 (Koutango), and 8.

According to the Brazilian Ministry of Health, WNV has been circulating in Brazil’s northeast region since 2014^6^, although there has been serological evidence for its circulation since 2004^3^. Here, we confirm the WNV circulation in Brazil, as we were able to isolate and acquire the complete genome sequence of a second isolate [(GenBank accession: MT905060 (JNESEq804 strain)]. Although NGS data was inconclusive to obtain the WNV complete genome sequencer from 827_01_CE, we also provided the evidence for WNV circulation in Ceará state as 827_01_CE WNV was independently confirmed by RT-qPCR in both LFDA/MG (Ct 32.1) and LVM-FMRP-USP (Ct 27.6), and confirmed by Sanger sequencing.

Our data demonstrate that WNV NY99 strain has been circulating silently in Brazil and in South America (SA) for about two decades and reinforces that WNV was probably introduced in the country many years before the first official report; probably between 2001 and 2005^5^. Up to this date, all sequenced SA WNV strains belong to the NY99 strain clade that was displaced by two other strains [WN02 (env-V159A) and SW03 (NS4a-A85T)] in the USA, between 2001 and 2003. Interestingly, after 2005, the NY99 strain completely disappeared from the USA^9^.

Despite the evidence that WNV has been circulating in Brazil for quite some time many questions still remain: (1) How was WNV introduced in Brazil and how has the NY99 signature been preserved until now?; (2) Why wasn’t it detected until 2014/2018?; (3) Why has no other strain (WN02/SW03) been introduced or detected?; (4) Which mosquitoes and birds species WNV is using to be amplified in the Brazilian territory?; (5) Is our major prevalent vector (*Culex quinquefasciatus*) capable of efficient transmission as *Culex* mosquitoes in the USA?; (6) If WNV has been circulating in Brazil for many years, why an outbreak of human infections has not occurred yet?; (7) Is this due to cross-protection with other endemic flavivirus (DENV, YFV, SLEV, or ZIKV)?

Most of these questions cannot be answered because the available data are not enough to draw any conclusion, but it is almost certain that WNV was probably introduced in Brazil by bird’s migration or by legal/illegal trade. Brazil is a tropical country and has one of the most diverse fauna in the world, and this could be a protective factor against an outbreak, reducing the viral activity^10^. To try to fill all these gaps, it is necessary to make the surveillance system sensible enough to identify suspicious cases of neurological disease in humans and animals, and to perform differential diagnoses for other virus species. Applying One Health concept and understanding that animals, in general, could be used as sentinels will provide more data to understand the epidemiology of WNV in the Americas, and help to detect its early circulation before it causes an explosive outbreak, as happened in the USA during 1999, with many human fatal cases.

## Ethics

Animal samples were collected by the Official Veterinary Service of each state and sent to the national reference laboratory, Federal Agricultural Defense Laboratory of Minas Gerais (LFDA/MG), from the Ministry of Agriculture, Livestock, and Supply (MAPA) to perform the diagnosis. Both animals died naturally. This study does not require approval by Ethics Committees according to Brazilian normative resolution n° 30, of February 2, 2016 of the National Animal Experimentation Control Council - CONCEA.

For IFA was performed with hyperimmune serum obtained by venipuncture from infected mice, the Ethics Committee on the Use of Animals (CEUA) from Ribeirão Preto Medical School - University of São Paulo approved the study under process number 214/2019.

## Author’s contributions

**MJLS**: Conceptualization, Methodology, Investigation, Data Curation, Writing - Original Draft, Writing - Review & Editing, Visualization, Project administration, Funding acquisition; **DMMJ**: Conceptualization, Software, Investigation, Resources, Writing - Original Draft, Writing - Review & Editing; **LACJ**: Visualization, Data Curation, Writing - Review & Editing; **AAF-J, MLN, MFC** and **EDLC**: Investigation, Resources; **VGF, FRS** and **EMQ-J**: Investigation; **AABC**: Supervision, Writing - Review & Editing; **BALF**: Conceptualization, Methodology, Resources, Writing - Review & Editing, Supervision, Project administration, Funding acquisition. All authors read, reviewed and approved the final version of the manuscript

## Acknowledgments

We thank Renato David Puga, MSc, for helping to analyze the sequence of 827_01_ES.

## References

1. Schweitzer BK, Chapman NM, Iwen PC. Overview of the Flaviviridae With an Emphasis on the Japanese Encephalitis Group Viruses. LabMedicine. 2009;40(8):493–9

2. Simmonds P, Becher B, Bukh J, Gould EA, Meyers G, Monath T, et al. ICTV Virus Taxonomy Profile: Flaviviridae. J Gen Virol. 2017;98:2–3.

3. Castro-Jorge LA, Siconelli MJL, Ribeiro BS, Moraes FM, Moraes JB, Agostinho MR, Klein TM, et al. West Nile virus infections are here! Are we prepared to face another flavivirus epidemic? Rev Soc Bras Med Trop. 2019;52:1–9

4. Lanciotti RS, Roehrig JT, Deubel V, Smith J, Parker M, Steele K, et al. Origin of the West Nile virus responsible for an outbreak of encephalitis in the northeastern United States. Science. 1999;286:2333–7.

5. Martins LC, Silva EVP, Casseb LMN, Silva SP, Cruz ACR, Pantoja JASP, et al. First isolation of West Nile virus in Brazil. Mem Inst Oswaldo Cruz. 2019;114

6. Vieira MACS, Romano APM, Borba AS, Silva EVP, Chiang JO, Eulálio KD, et al. West Nile Virus Encephalitis: The First Human Case Recorded in Brazil. Am J Trop Med Hyg. 2015;93(2):377–9.

7. Eiden M, Vina-Rodriguez A, Hoffmann B, Ziegler U, Groschup MH. Two new real-time quantitative reverse transcription polymerase chain reaction assays with unique target sites for the specific and sensitive detection of lineages 1 and 2 West Nile virus strains. J Vet Diagn Invest. 2010;22:748–53.

8. OIE. World Organization for Animal Health. West Nile Fever. In: OIE Terrestrial Manual, 2018.

9. Hadfield J, Brito AF, Swetnam DM, Vogels CBF, Tokarz RE, Andersen KG, et al. Twenty years of West Nile virus spread and evolution in the Americas visualized by Nextstrain. Plos Pathog. 2019;15:1–18.

10. Diaz LA, Flores FS, Quaglia A, Contigiani MS. Intertwined arbovirus transmission activity: reassessing the transmission cycle paradigm. Front Physiol. 2013;3(493):1–7.

